# Primary platinum resistance and immune exclusion in ovarian carcinomas with high expression of the homologous recombination mediator RAD51

**DOI:** 10.1101/2020.06.09.137612

**Authors:** Michal M. Hoppe, Tuan Zea Tan, Stefanus Lie, Sherlly Lim, Joe PS Yeong, Joanna D. Wardyn, Patrick Jaynes, Diana GZ Lim, Brendan NK Pang, Anthony Karnezis, Derek S. Chiu, Samuel Leung, David G. Huntsman, Anna S. Sedukhina, Ko Sato, Stan Kaye, Hyungwon Choi, Naina Patel, Robert Brown, Jason J. Pitt, David SP Tan, Anand D. Jeyasekharan

**Affiliations:** Cancer Science Institute of Singapore, National University of Singapore, Singapore; Department of Pathology, National University Hospital, Singapore; British Columbia Cancer Agency, Vancouver, Canada; Department of Pharmacogenomics, St. Marianna University, Kawasaki, Japan; Cancer Research UK Clinical Trials Unit, University of Glasgow, UK; Scottish Gynaecological Cancer Trials Group (SGCTG), UK; Saw Swee Hock School of Public Health, National University of Singapore, Singapore; Division of Cancer and Ovarian Cancer Action Research Centre, Department of Surgery and Cancer, Imperial College London, London, UK; Department of Haematology-Oncology, National University Hospital, Singapore

**Keywords:** Quantitative imaging, digital pathology, mIHC, RAD51, Ovarian Cancer, HRD, immune exclusion

## Abstract

**Background:** Homologous recombination deficiency (HRD) in ovarian cancer confers increased sensitivity to Poly-ADP-Ribose-Polymerase (PARP) inhibitors and platinum. The homologous recombination (HR) mediator RAD51, however, is commonly overexpressed, potentially driving unregulated HR. Due to non-quantitative measurements and heterogeneous cohorts, the clinical relevance of RAD51 expression in ovarian cancer is unclear.

**Methods:** Fluorescent immunohistochemistry and multispectral imaging were used to quantitate RAD51 expression, in relation to other markers, in independent cohorts of ovarian carcinomas from British Columbia Cancer (BCC *n*=284) and the phase III SCOTROC4 trial (*n*=268). Independent cohorts (TCGA *n*=566, GSE9891 *n*=267, GSE26712 *n*=185 and GSE3149 *n*=146) were used for mRNA expression and immune infiltration analyses.

**Results:** RAD51-High tumours had shorter progression-free and overall survival compared to RAD51-Low cases in both BCC and SCOTROC4. The negative prognostic significance of high RAD51 was primarily evident in cases classified “HRD negative” by the Myriad genomic scar assay. Unexpectedly, overexpression of RAD51 in ovarian cancer cell lines did not affect sensitivity to platinum or PARP inhibitors, but modified the expression of immune-regulatory genes. Accordingly, tumours with high *RAD51* mRNA showed consistent changes in immunomodulatory transcripts across four independent ovarian cancer cohorts. In-situ multiplexed imaging confirmed that high RAD51 tumours correlated with tumour exclusion of cytotoxic-T-cells, possibly explaining their poor outcomes after chemotherapy.

**Conclusions:** High RAD51 expression, in conjunction with a HRD score <42, identifies a subgroup of EOC cases with platinum resistance and an immune excluded tumour microenvironment. Tumours with high RAD51 may require alternate or additional adjuvant therapeutic strategies to overcome immune exclusion.

**Statement of Translational Relevance:** Homologous recombination deficiency (HRD) in ovarian cancer confers increased sensitivity to Poly-ADP-Ribose-Polymerase (PARP) inhibitors and platinum. Currently, methods to identify patients who do poorly on first-line platinum based chemotherapy in ovarian cancer represent an unmet clinical need. In this report we show that RAD51 expression can identify patients with primary platinum resistance in two large cohorts, including a unique phase-III trial of carboplatin as adjuvant monotherapy. The clinical relevance of a positive genomic scar HRD score, a biomarker for assessing HRD status, in predicting benefit to PARP inhibitors has been established following recent phase 3 trials in first line and 2^nd^ line maintenance treatment of ovarian cancer. Our findings suggest that HRD negative ovarian tumours may be further stratified by RAD51 expression status in the context of platinum sensitivity, and may serve to guide future trials of HRD targeted agents.

## Introduction

Homologous recombination (1,2) (HR) is a DNA repair pathway of clinical interest due to the enhanced sensitivity of HR-deficient (HRD) cells and tumours to platinum chemotherapy and PARP inhibitors (PARPi) (3-6). However, the opposite of HRD i.e. hyper-recombination or a hyperactive HR state, also occurs in cancer (7). For example, in Bloom Syndrome (8) mutations in the *BLM* DNA helicase gene lead to increased illegitimate recombination events. However, the clinical relevance of markers that indicate hyperactive HR in somatic cancers is unclear. One such potential marker is RAD51, the vertebrate homolog of bacterial RecA and a central, conserved part of the HR reaction. RAD51 forms a nucleoprotein filament on single-strand DNA which mediates the process of strand invasion into the homologous sister chromatid during HR. In-vitro, overexpression of RAD51 leads to unregulated, hyperactive HR (9-13). Depletion or mutation of RAD51 is lethal, consistent with the essential role of RAD51 mediated strand-exchange in resolving abnormal replication intermediates (14,15). As such, unlike other HR components, RAD51 is not commonly mutated in cancer. Conversely, it is often upregulated and is associated with poor survival (16-21). Evaluating the clinical significance of RAD51 overexpression, especially in relation to other HR defects, has been hampered by the lack of quantitative tools for proteomics in-situ. In this paper we utilize a quantitative platform based on multispectral imaging and automated analysis to evaluate baseline RAD51 protein levels in formalin fixed paraffin embedded (FFPE) tissue. For these studies we focus on epithelial ovarian cancer (EOC), a tumour type where deregulation of the HR pathway is common (22). High-grade serous ovarian carcinomas (HGSOC) are the most common and aggressive subtype of EOC, and we first evaluated RAD51 in a cohort of HGSOC. We show that tumours with high basal (pre-treatment) RAD51 protein expression have poorer survival outcomes. Platinum-based chemotherapy is the mainstay of the management of EOC but it is typically combined with paclitaxel, the sensitivity to which is not associated with HRD, making it challenging to dissect the contribution of HR to survival. We therefore confirmed our findings in an independent cohort of the SCOTROC4 clinical trial. SCOTROC4 was a phase III clinical trial performed to assess two different dosing schedules of single agent carboplatin in EOC (23). While the trial showed no difference between the two arms, it represents a unique cohort of single agent platinum treated ovarian cancer with well-annotated survival data and HRD scores (24). Finally, we use a combination of in-vitro experiments and RNA expression data from four independent ovarian cancer datasets to demonstrate that the effect of RAD51 overexpression on survival is potentially mediated through an altered immune microenvironment in RAD51-High EOC.

## Materials and Methods

### Patients and treatment

EOC FFPE samples to optimize RAD51 fluorescent immunohistochemistry (fIHC) were obtained from National University Hospital of Singapore (NUH cohort, *n=*52). A cohort of HGSOC was obtained from British Columbia Cancer (BCC TMA, *n*=308, SuppTable-1); samples of EOC from the SCOTROC4 clinical trial (*n=*309, SuppTable-2), chosen based on sample availability and divided equally between both study arms(23), were used as a tissue microarray (TMA). HRD score from the Myriad genomic scar assay was available for 240/309 cases on this TMA(24). Gene expression analysis, was performed on HGSOC and high grade endometrioid samples from the TCGA (*n=*566)(22), Australian Ovarian Cancer Study (AOCS, GSE9891; *n*=267)(25), Massachusetts General Hospital (MGH, GSE26712; *n*=185)(26) and Duke University Hospital (Duke, GSE3149; *n*=146)(27), which were extracted from CSIOVDB(28). Tumour and data collection was approved by local ethics committees for all the above cohorts.

### Fluorescent immunohistochemistry (fIHC) and gene expression

Multiplexed fIHC was performed on FFPE samples to assess protein expression using an Opal 7-Color Kit, and imaged/analysed using the Vectra 2 System (PerkinElmer Inc., Waltham, MA, USA). The RAD51 Vectra score (RAD51_Vs_) measures mean nuclear fluorescent intensity per tumour expressed in normalized counts. Antibodies used are in Supplementary table 3. The results of this study are reported in concordance with REMARK guidelines (29). Complete details of the fIHC and gene expression analysis methods are presented in the supplementary information.

### Statistical methods

For categorical analyses, quantitative scores were divided into three groups accordingly - first quartile (Q1), interquartile range (IQR) and fourth quartile (Q4). For the BCC cohort, progression free survival (PFS) and overall survival (OS) were defined as time from the date of diagnosis to progression or death respectively. For the SCOTROC4 data, PFS and OS were defined as time from the date of randomization to progression or death (PFS) or death (OS) and estimated using the Kaplan–Meier method and tumour progression was determined according to RECIST version 1.0 criteria. CT scans were carried out at baseline and after six cycles of treatment and if CA125 rose or clinical progression was suspected (23). Kapan-Meier curves are shown for Q4 and Q1. Cox proportional hazards (Cox PH) models were calculated using IBM SPSS Statistics 23 software; all variables satisfied the proportional hazards assumption and all clinically relevant clinicopathological variables were included in the multivariate models. Other statistical tests and graphs were generated using GraphPad Prism 8 software. Statistical tests were two-sided and *p*<0.050 was considered as statistically significant in individual testing; Bonferroni correction was applied to pairwise comparisons when more than two groups were present.

## Results

### Baseline RAD51 protein expression is associated with poorer survival in ovarian cancer

Quantitative analysis of protein expression by fluorescent immunohistochemistry (fIHC) is critically dependent on antibody specificity. We validated clone EPR4030(3) (Abcam) to reliably detect baseline levels of RAD51 in FFPE samples using RAD51 siRNA treated cell-blocks (SuppFig.1A-C). Automated Vectra scoring of multiplexed RAD51 fIHC on a cohort of EOC cases (*n*=52) showed strong concordance with the H-Score defined visually by two independent pathologists (SuppFig.1D). We then applied this optimized protocol for staining, imaging and scoring to estimate RAD51 protein expression in ovarian cancer to generate a RAD51 Vectra score (RAD51_Vs_) per case.

Histological subtypes of ovarian cancers have distinct biological and clinical characteristics. We therefore evaluated RAD51 expression in a cohort of HGSOC, the most common subtype. The RAD51_Vs_ in the BCC cohort of HGSOC followed a normal distribution (Fig.1A). To ascertain the clinical significance of high basal RAD51 expression, we performed a categorical survival analysis on the BCC cohort according to the RAD51_Vs_, by dividing the cohort into three subsets. RAD51-Low cases were defined as those within the first quartile (Q1), RAD51-High cases as cases within fourth quartile (Q4), and RAD51-IQR as cases within the interquartile range (IQR, quartiles 2+3) (Fig. 1B). In a Kaplan-Meier survival analysis, we noted an association of RAD51_Vs_ with survival (median PFS 1.1 vs. 1.7 years, p<0.001; median OS 2.7 vs. 3.8 years, p=0.001, for RAD51-High and -Low respectively; log-rank test) (Fig.1C). RAD51-High cases also showed higher likelihood of progression than RAD51-Low cases (odds ratio (OR) for progression at 12 months 2.5, *p*=0.018; at 24 months 3.9, *p*=0.002) (Fig.1D).

**Figure 1.**
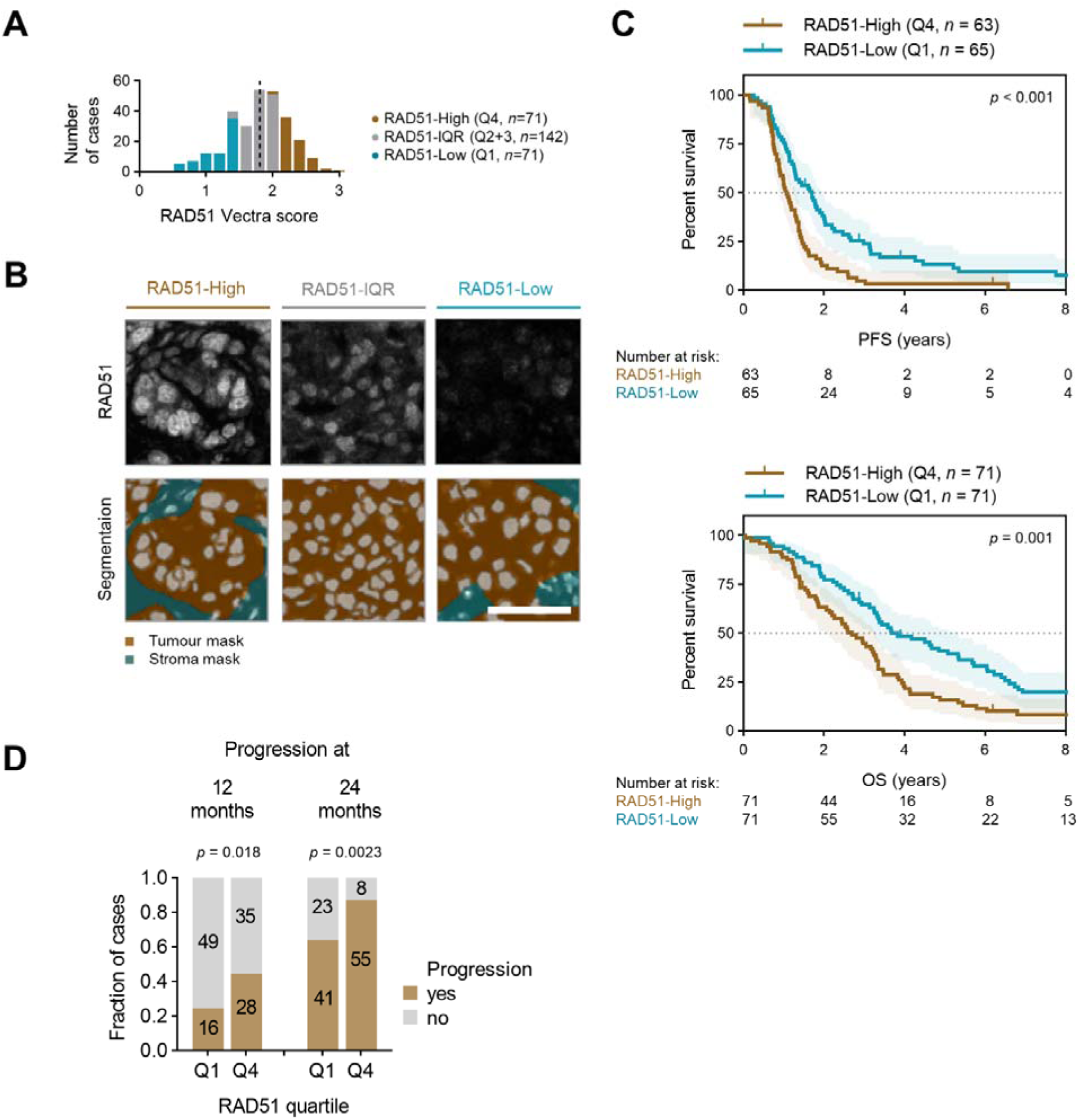
RAD51 protein expression in EOC. **A**, Distribution of RAD51 Vectra score (RAD51_Vs_) in the BCC cohort. The cohort is divided into RAD51-Low expressing cases (first quartile, Q1 – blue), intermediate cases (interquartile range, IQR – grey) and RAD51-High expressing cases (fourth quartile, Q4 – blue). **B**, Example unmixed RAD51 staining images *(top panel*) with respective tissue segmentation and cell segmentation mask. The tumour mask is based on EPCAM expression positivity. Scale bar is 50μm. **C**, Survival analysis of the BCC cohort. Kaplan-Meier plots for progression-free survival (PFS) (*top*) and overall survival (OS) (*bottom*) stratified according to fourth quartile (Q4) and first quartile (Q1) of RAD51_Vs_. Log-rank test. **D**, Number of cases with progression at 12 and 24 months. Chi-square test.

### RAD51-High tumours show poor survival independent of proliferation status

As RAD51 expression is linked to proliferation through common regulatory pathways (30), a possible explanation for the poorer survival could be increased proliferation in RAD51-High tumours. We therefore measured Ki67 expression by fIHC and assigned a percentage-positivity score for the BCC cohort cases. RAD51_Vs_ correlated weakly with Ki67 extent (*r*=0.251, *p*<0.001, Spearman correlation) (Fig.2A). We performed an fIHC co-staining with Geminin (S/G2 cell cycle-phase marker) along with RAD51 and note that at a single-cell resolution, RAD51 expression is indeed higher in cells in the S/G2 cell cycle-phase (Fig.2B). Nonetheless, there was RAD51 expression noted in quiescent and G1 cells, which was also elevated in RAD51-High tumours (Fig.2B). These findings suggest the presence of a cell-cycle/proliferation independent regulation of RAD51 expression in tumours. Accordingly, the proliferation status of the tumour (i.e. extent of Ki67 positivity) was not associated with survival outcomes (Fig.2C). Furthermore, in a multivariate cox proportional hazards (Cox PH) model adjusting for Ki67 extent, age and stage, RAD51_Vs_ as a continuous variable remained a statistically significant independent predictor of PFS in HGSOC (HR 1.4 per unit of RAD51_Vs_, 95%CI 1.0 to 1.9, *p*=0.025; Cox PH). Comparable results were obtained for OS (HR 1.3 per unit of RAD51_Vs_, 95%CI 0.98 to 1.9, *p*=0.066; Cox PH) (Table 1).

**Figure 2.**
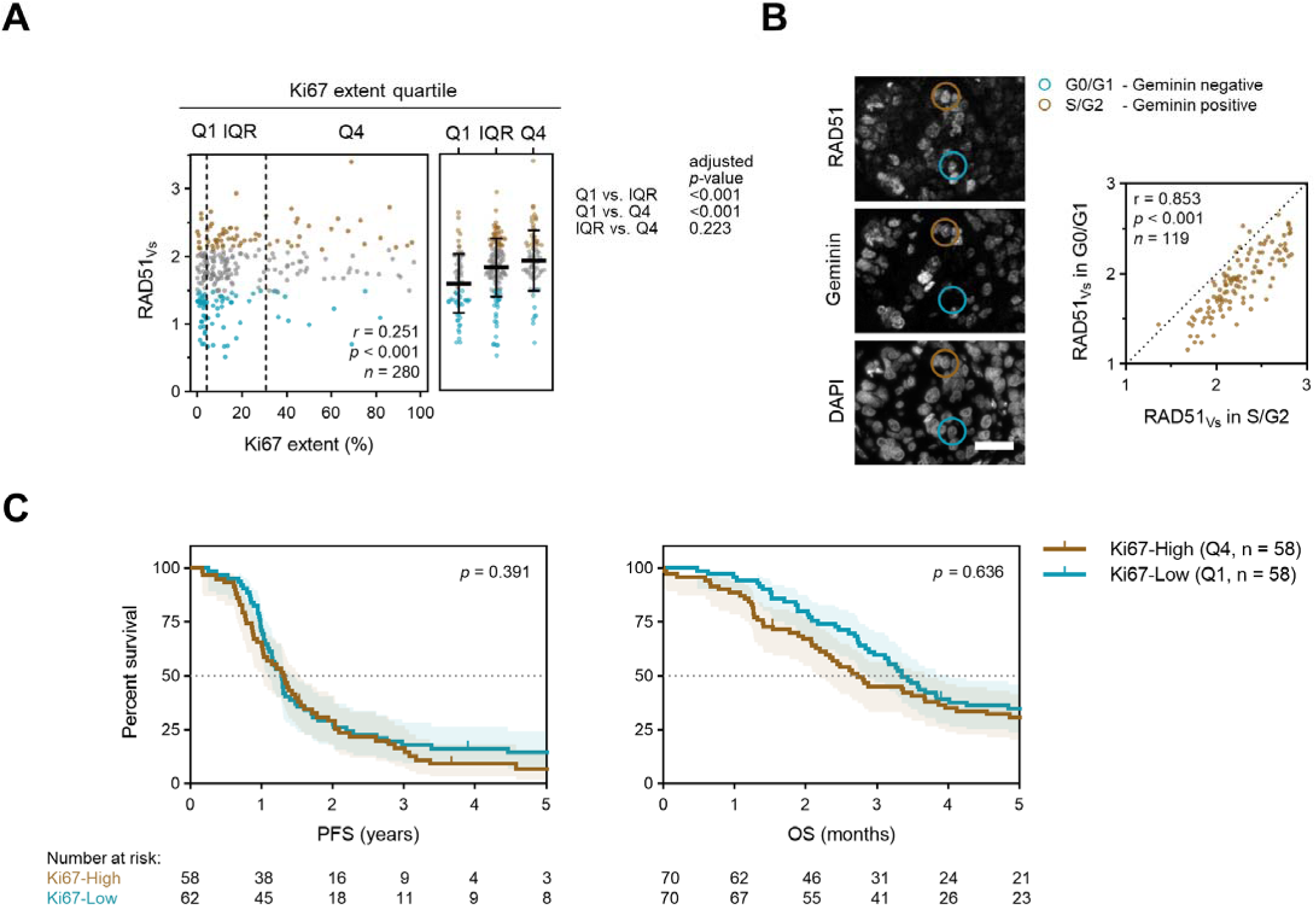
Proliferative capacity is not associated with survival in EOC. **A**, Correlation of Ki67 extent and RAD51 Vectra score (RAD51_Vs_) in the BCC cohort. Spearman correlation (*left*) and one-way ANOVA with Bonferroni correction (*right*). Mean with standard deviation. **B**, RAD51 and Geminin co-staining (*left*) and quantitation of RAD51_Vs_ in the G0/G1 and S/G2 subpopulations based on Geminin positivity. Scale bar is 50μm. Spearman correlation. **C**, Kaplan-Meier plots for PFS (*left*) and OS (*right*) stratified according to Ki67 extent quartile. Q – quartile. Log-rank test.

**Table-1.**
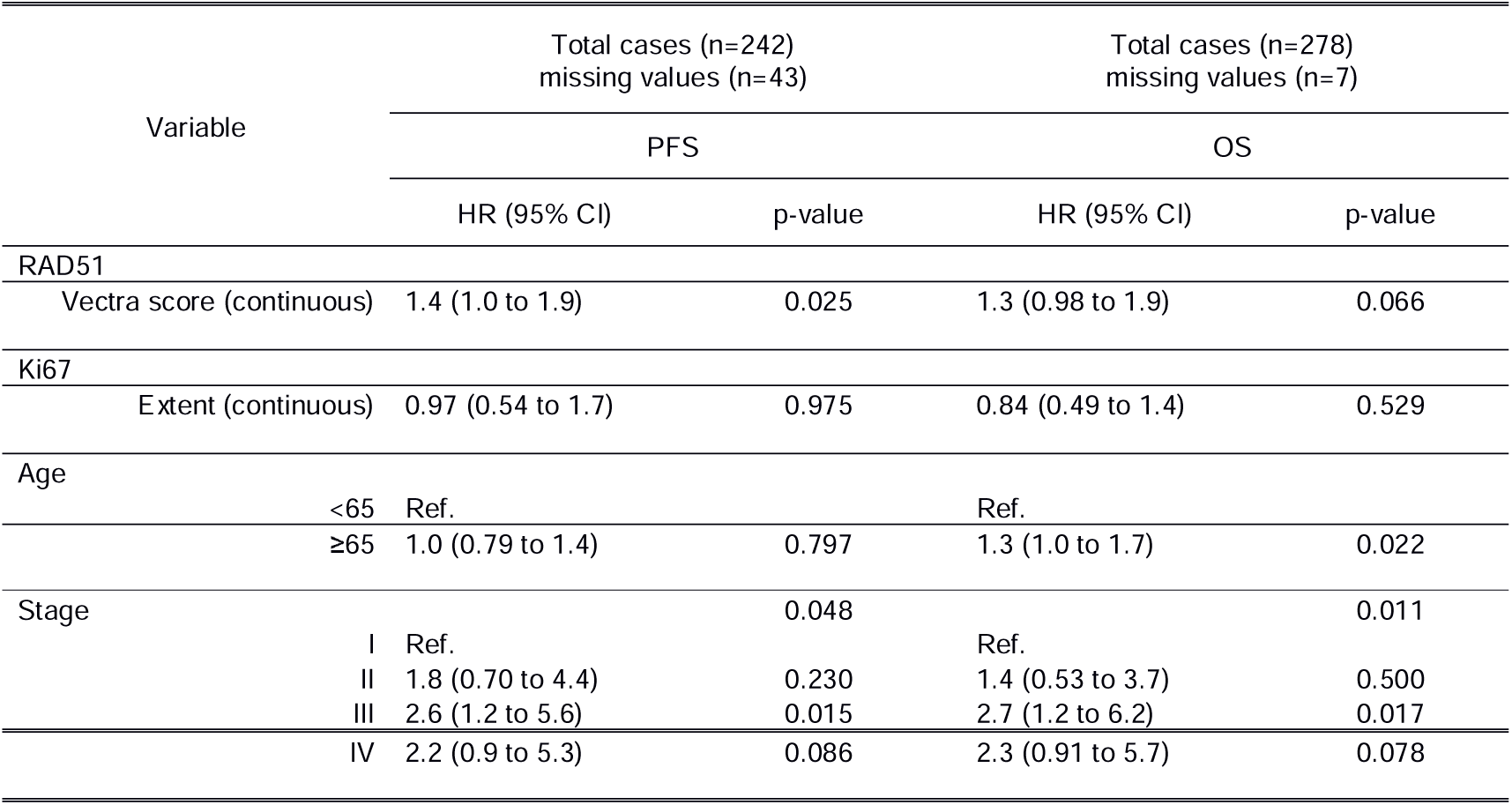
Multivariate analysis of continuous RAD51 Vectra score and Ki67 extent as a predictor of PFS and OS in the BCC cohort of HGSOC (Cox proportional hazards model).

### Independent validation of RAD51 Vectra score with outcomes in EOC from SCOTROC4

RAD51-dependent recombination is a pathway for the repair of DNA damage caused by platinum chemotherapy. To negate potential confounding effects of paclitaxel on survival outcomes, which is typically administered in combination with carboplatin in the first line for ovarian cancer, we utilized the unique EOC SCOTROC4 trial cohort, which were treated with carboplatin monotherapy. RAD51 protein expression followed a normal distribution within this cohort as well (Fig.3A), and a relationship with proliferation status was noted (*r*=0.468, *p*<0.001, Spearman correlation) (SuppFig.2A). Again, we observe a subset of RAD51-High cases with low Ki67 extent. RAD51-High patients showed poorer PFS and OS after platinum monotherapy in comparison to RAD51-Low cases (median PFS 0.99 vs. 1.2 years, *p*=0.013; median OS 2.3 vs 4.0 years, *p*=0.002, for RAD51-High and -Low respectively; log-rank) (Fig.3B). Data on adjuvant treatment end dates from the SCOTROC4 trial for were not available for us to calculate the platinum-free interval (PFI) from completion of treatment to time of progression, however, all patients in SCOTROC4 commenced treatment within 8 weeks of surgery as per protocol and ∼80% of patients completed 6 cycles of 3 weekly paclitaxel (23). We therefore used the PFS at 12 months (calculated from time of randomisation) as a surrogate for a shorter PFI and hence platinum resistance. RAD51-High cases were more likely to relapse within 12 months compared with RAD51-Low cases (57% vs 38% 12 month relapse rate respectively, p=0.039) (Fig.3C), thus suggesting a higher risk of primary platinum resistance in RAD51-High tumours. At the 24 month PFS timepoint, the difference was even more marked with a 91% vs 60% relapse rate for RAD51-High versus RAD51-Low tumours respectively (*p*<0.001) (Fig.3C). In this cohort, there was an association of Ki67 with poorer PFS in a univariate analysis, but no effect of Ki67 on survival was observed in OS analysis (SuppFig.2B). We then performed a Cox PH multivariate analysis of both continuous linear RAD51_Vs_ and Ki67 extent taking histology into account; along with age, stage, grade, performance status, bulk of residual disease and HRD status (Table 2). RAD51_Vs_ as a continuous variable was not an independent predictor of PFS, likely due to its correlation with tumour stage in this cohort (HR for PFS 1.2 per unit of RAD51_Vs_, 95%CI 0.97 to 1.5, *p*=0.104; Cox PH), however it remained an independent statistically significant predictor of OS (HR for OS 1.4 per unit of RAD51_Vs_, 95%CI 1.1 to 1.9, *p*=0.007; Cox PH).

**Figure 3.**
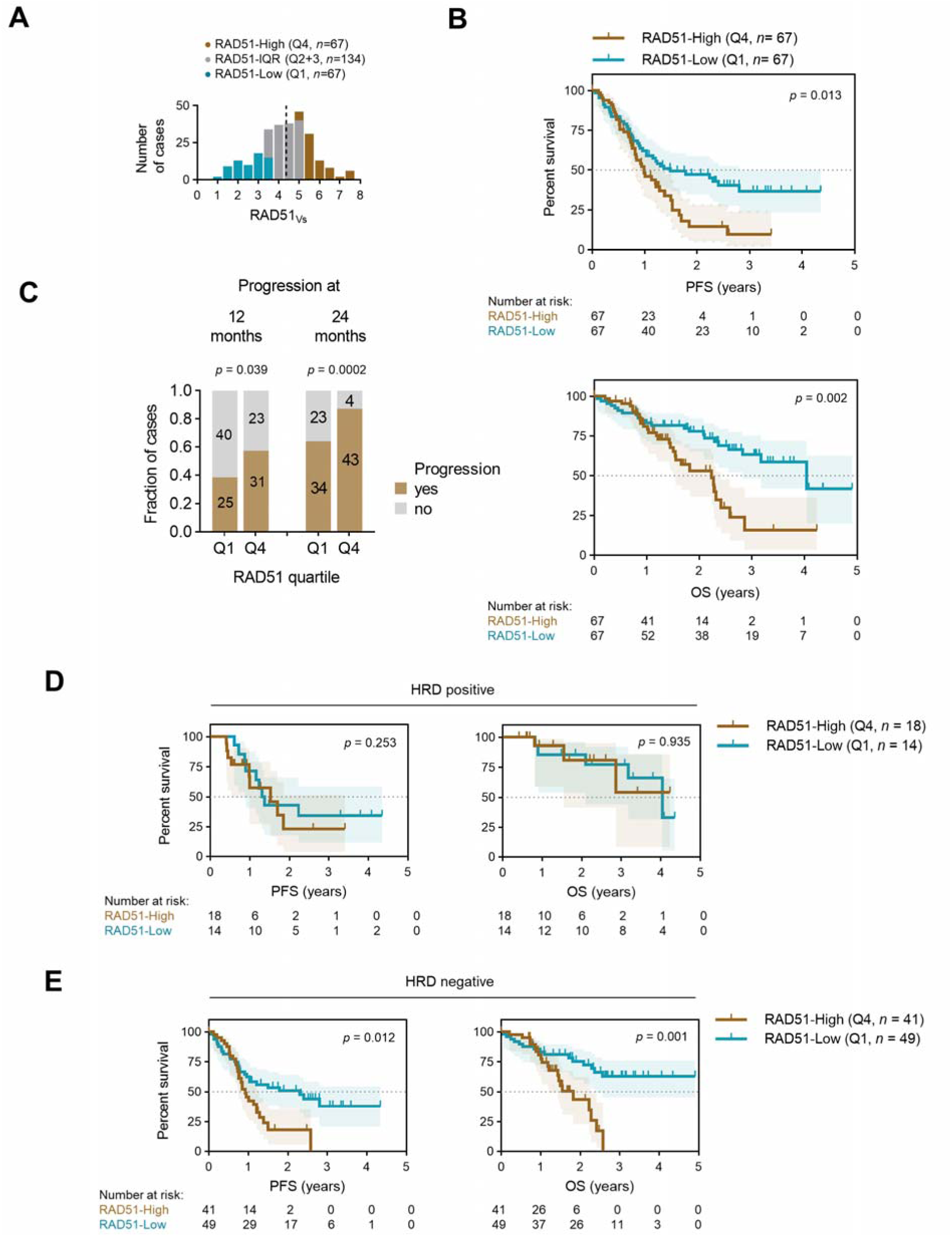
Synergy between RAD51_Vs_ and HRD score. **A**, Distribution of RAD51 Vectra score (RAD51_Vs_) in the SCOTROC4 cohort. **B**, Survival analysis of the SCOTROC4 cohort. Kaplan-Meier plots for PFS (*left*) and OS (*right*) stratified according to quartiles of RAD51_Vs_. **C**, Number of cases with progression at 12 and 24 months. Chi-square test. **D**, Survival analysis of HRD positive patients according to quartile of RAD51_Vs_. **E**, Survival analysis of HRD negative patients. Log-rank test.

**Table-2.**
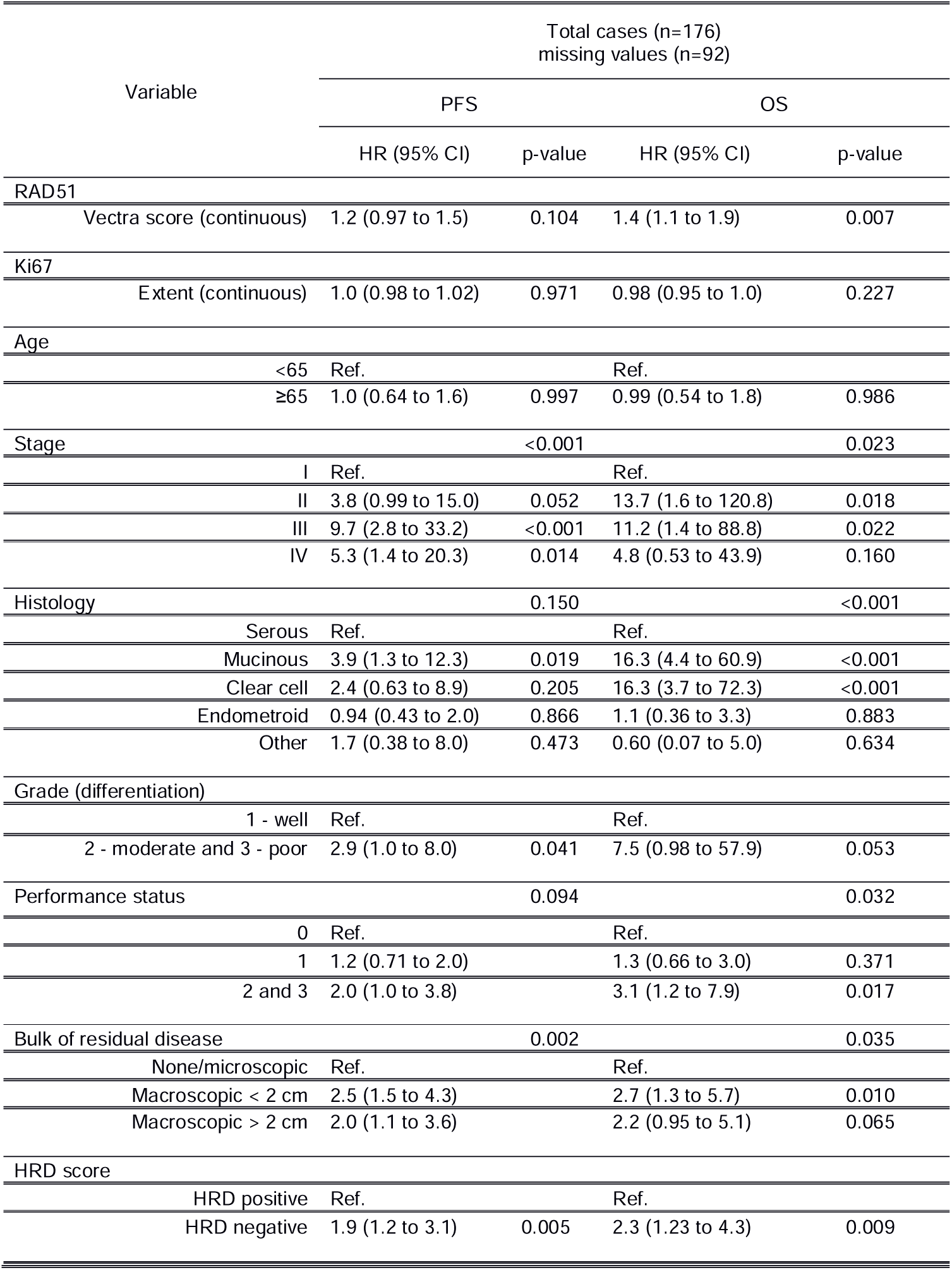
Multivariate analysis of continuous RAD51 Vectra score and Ki67 extent as a predictor of PFS and OS in the SCOTROC4 cohort (Cox proportional hazards model).

### RAD51 Vectra score predicts poor outcomes in HRD negative EOC patients

Overexpression of RAD51 may be a possible mechanism to overcome HRD (31). However, we did not observe any difference in RAD51_Vs_ between *BRCA* mutant (typically HRD positive) and non-mutant cases in terms of RAD51 expression (SuppFig. 3A). HRD can also result from non-*BRCA* genetic or epigenetic alterations, which are captured in genomic scar assays. The Myriad Genetics HRD score was available for 240 patients in the SCOTROC4 cohort and 67 (27.9%) were defined as HRD positive (based on a validated cut-off of >42) (32,33), with longer PFS and OS on platinum treatment compared to HRD negative patients (24) (reanalysed in SuppFig.3B). HRD positivity was observed in 28/31 (90.3%) of *BRCA-* mutant. cases in this cohort, confirming the sensitivity of the assay. Similar to *BRCA* mutations, absolute HRD scores did not associate with RAD51_Vs_ (SuppFig. 3C). Finally, we did not observe any association between RAD51_Vs_ and DNA damage response markers measured by fIHC (SuppFig. 3D). Together, these results suggest that RAD51 expression *in vivo* is not driven by underlying HRD or chronic activation of the DNA damage response.

To explore the clinical relevance of RAD51 expression in the context of HRD, we performed a subset survival analysis of HRD positive and negative tumours in the SCOTROC4 cohort. Within the HRD positive group, the RAD51_Vs_ did not stratify patients for survival (Fig.3D). However, we observed a clear association of the RAD51_Vs_ with survival within the HRD negative subgroup; RAD51-Low cases showed an increase in PFS and OS compared to RAD51-High cases (median PFS 0.92 vs. 2.3 years, *p*=0.012; median OS 1.8 vs. not reached, *p*=0.001, for RAD51-High, and -Low respectively; log-rank) (Fig.3E). The association of RAD51 with OS remained significant in this HRD negative group using a multivariate Cox PH analysis (SuppTable-4). A similar trend was noted in the HRD negative HGSOC subset (median OS 1.6 vs. not reached for RAD51-High and -Low respectively, *p*=0.035; log-rank) (SuppFig.3E). The 12 month PFS rate was also higher in the RAD51-High vs RAD51-Low cases (59% versus 38%, Chi-squared *p*=0.068, data no shown) in the SCOTROC 4 HRD negative subgroup. These results suggest that RAD51 expression is most relevant in predicting for platinum resistance within the subgroup of EOC cases defined by the genomic scar assay as HRD negative. These two assays utilized together therefore identify a subset of cases that are least likely to respond to platinum treatment.

### RAD51-High tumours constitute a novel subset of ovarian cancer with a distinct immunological profile

To test whether high expression levels of RAD51 may cause resistance to platinum therapy in-vitro, we developed a RAD51 overexpression model in HGSOC cell lines (34) (SuppFig.4A-B). However, we did not observe resistance to platinum (Fig.4A) nor to the PARPi olaparib (SuppFig.5) upon RAD51 overexpression in-vitro. Since RAD51 expression did not associate with *BRCA* mutations, HRD or activation of the DDR, we sought to define the molecular characteristics of RAD51-High tumours to explain the discordance between the clinical findings and in-vitro findings with overexpression of RAD51. We performed a whole transcriptomic analysis between RAD51 overexpression and control HGSOC cell lines. A gene set enrichment analysis (GSEA) showed that RAD51-overexpression cell lines showed enrichment in the expression of genes related to regulation of T-cell and B-cell mediated immunity (Fig.4B). To substantiate these findings in clinical samples, we investigated differential gene expression of a curated set of immunity-related genes (35) (SuppTable-5) between RAD51-High and -Low tumours, which were defined by the upper (Q4) and lower (Q1) *RAD51* expression quartiles, respectively, in four independent cohorts of ovarian cancer: TCGA, AOCS, MGH and Duke cohorts (Fig.4C, as well as SuppFig.6). Remarkably we noted almost identical findings in all these four independent cohorts; RAD51-High tumours showed a distinct increase in expression of specific immune related genes. The highest association was noted for a cluster of immune genes which also link to proliferation-namely *TTK, PBK, CDK1* and *BRIC5*. However, several other *bona fide* immune markers such as *CXCL10, MICB, TFRC, CD47* and *ISG15* were also consistently high in RAD51-High tumours across all these clinical cohorts. In terms of immune processes, genes annotated Th1 orientation, and antigen presentation were enriched in RAD51-High tumours, while genes annotated in the complement were low in these tumours (Fig.4D).

**Figure 4.**
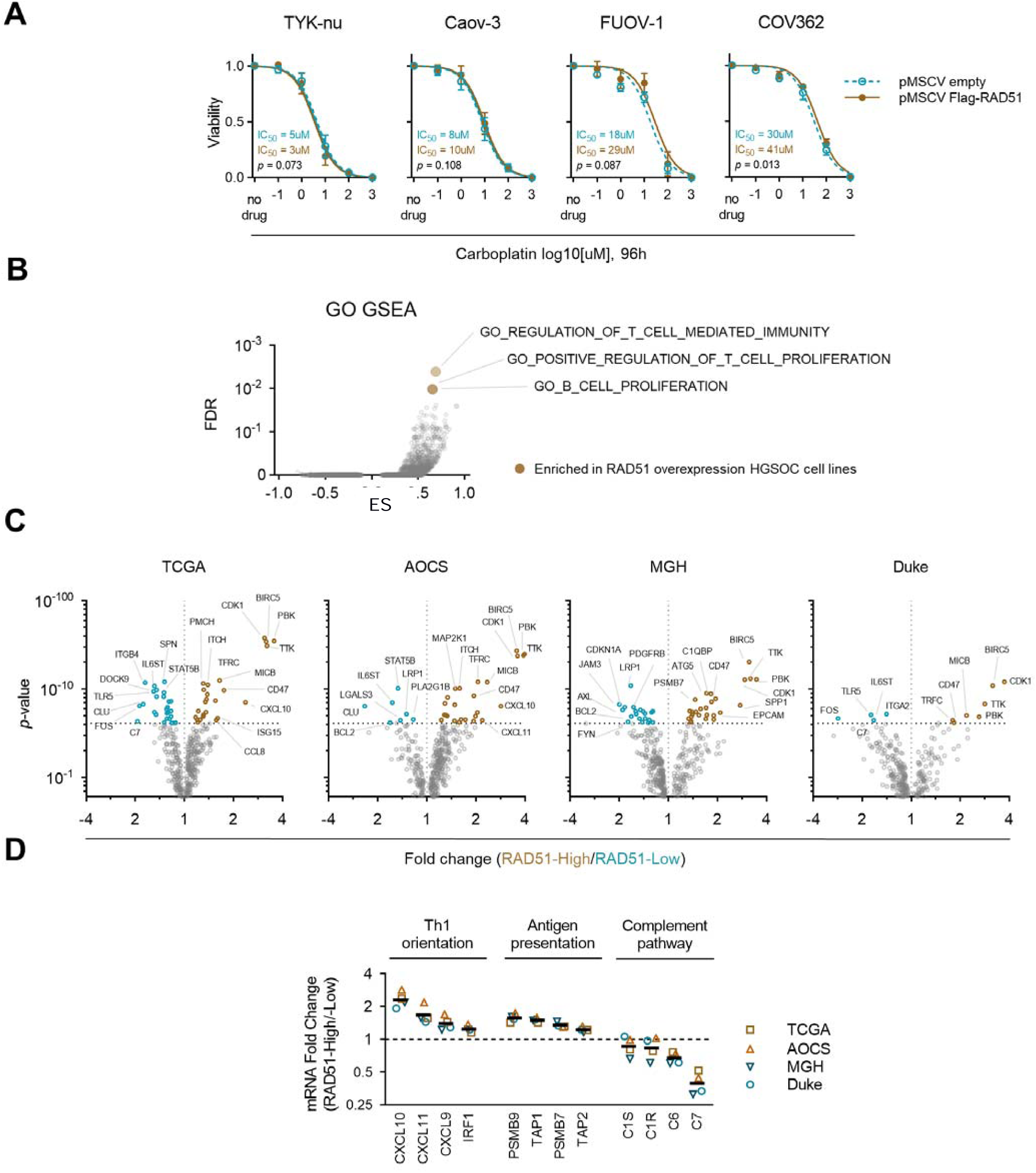
RAD51-High cases display an altered immune-microenvironment. **A**, *In vitro* cell survival assay of HGSOC cell lines upon RAD51 overexpression. At least three biological replicates per point. Extra-sum-of-squares F test. **B**, Gene Set Enrichment analysis of RAD51-overexpression and control HGSOC cell-lines (*n*=4). GO – Gene Ontology, ES – enrichment score, FDR – false-discovery rate. **C**, Differential gene expression analysis of immune genes between RAD51-High (Q4) and –Low (Q1) across four independent cohorts. **D**, Top consistently enriched genes with respective immune category.

### RAD51 overexpression is associated with cytotoxic T-cell exclusion *<u>in vivo</u>*

As the expression of several immune related genes was distinct in high-RAD51 tumours, we used spectral imaging to evaluate immune-infiltration in these tumours. To compare the immune microenvironment of RAD51-High and -Low tumours at a single-cell resolution *in situ*, we performed immune profiling of the BCC cohort by multiplexed fIHC of the markers CD3, CD8, FOXP3 and CD163, along with cytokeratin as a tumour mask (SuppFig.7). This panel allows the evaluation of immune infiltration by cytotoxic and regulatory T-cells, as well as pro-tumour macrophages, separately in the stroma and the tumour compartments. We observed an exclusion of CD3+/CD8+ cytotoxic T-cells in RAD51-High tumours (Fig.5A-B) with only weak changes in CD3+/FOXP3+ regulatory T-cell or CD163+ macrophage infiltration (SuppFig.8). Interestingly, the immune exclusion phenotype is primarily noted in tumours that are wild-type for the *BRCA* genes (Fig.5C), consistent with the survival significance of high RAD51 in HRD negative tumours. Taken together our findings suggest a possibility that high RAD51 expression constitutes a novel cell-intrinsic modulator of the tumour microenvironment, which leads to an immune excluded phenotype and poor survival.

**Figure 5.**
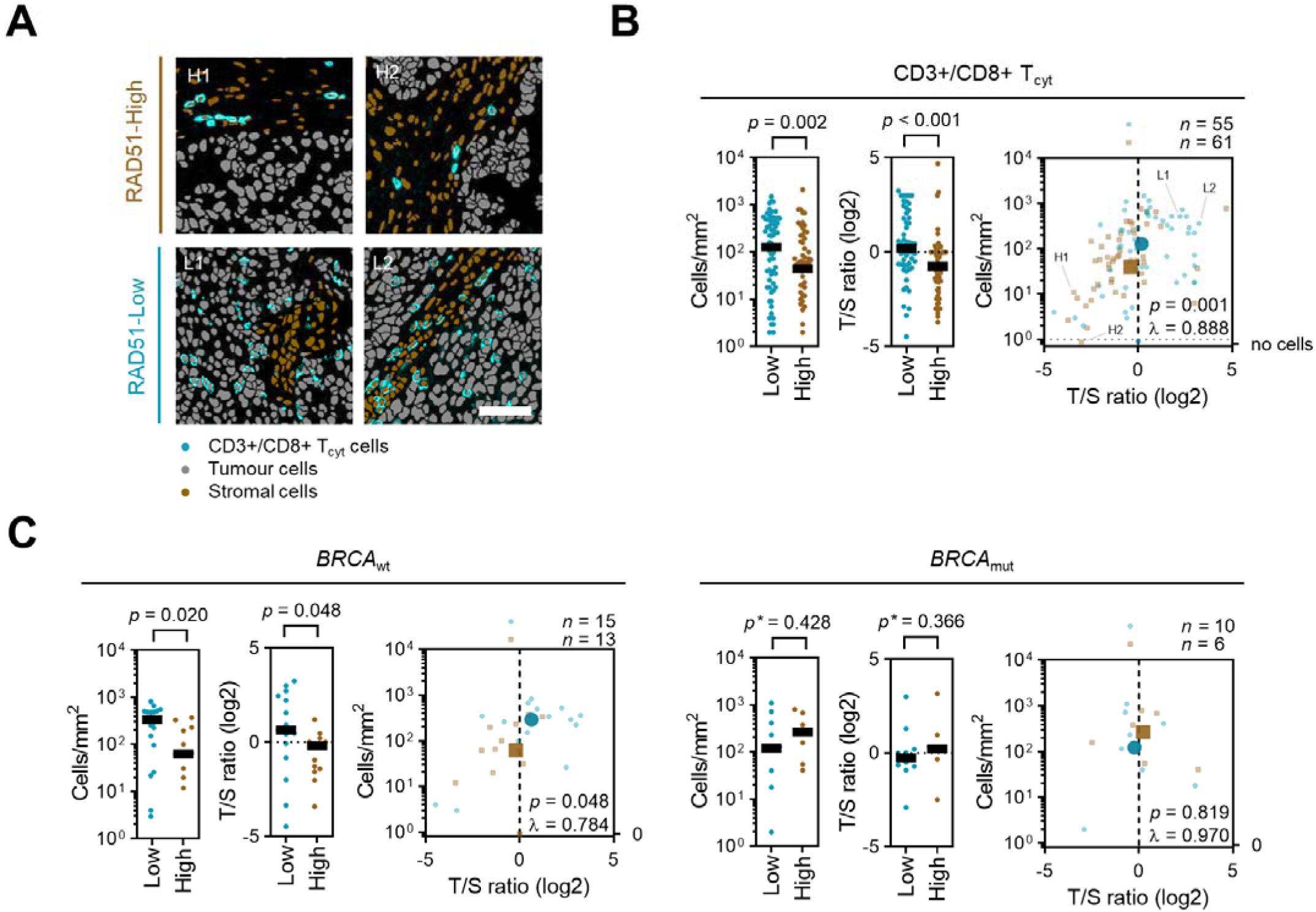
RAD51-High *BRCA* wild-type tumours show an impaired infiltration of cytotoxic T-cells. **A**, Example unmixed fIHC monochorme images of CD8, cytotoxic T-cell marker, overlaid on a tissue region-annotated cell segmentation mask of RAD51-High and -Low cases. All CD8-positive cells are also CD3-positive (not shown). Scale bar is 50μm. **B**, Cytotoxic T-cell infiltration analysis in the BCC cohort. Absolute tumour cytotoxic T-cell infiltration numbers and tumour/stroma (T/S) cytotoxic T-cell number ratio in RAD51-High and -Low cases *(left*). Bar is median. Mann-Whitney test. Spatial relationship of these two measurements is shown (*right*), centroids of both populations are represented as bold symbols. Example cases form (A) are marked on the graph. One-way MANOVA. **C**, Subset analysis as in (B) stratified according to *BRCA* mutation status.

## Discussion

In this study, we use cutting-edge techniques for the *in situ* quantitation of protein expression in clinical samples to evaluate RAD51, a key regulator of the homologous recombination pathway. RAD51 forms discrete nuclear foci upon activation of HR, and this is a widely used measure of recombination proficiency *in vitro*. The RAD51 foci counting assay has been evaluated in patient derived material and ex-vivo systems (36-39). However, the foci assay is not scalable, as automated and quantitative analysis of foci counts is challenging in FFPE material and highly reliant on sample preparation and microscopy setup. Aside from its phenotype of foci formation, RAD51 is also frequently overexpressed in cancer, the clinical relevance of which is not clearly established. In this paper, we have focused on the mean nuclear intensity (Vectra score) of RAD51 to measure expression of the protein in tumours, due to its feasibility for automated quantitation and scalability in large datasets.

We used multispectral imaging to study RAD51 in two independent cohorts of ovarian cancer. High RAD51 expression in both the BCC and SCOTROC4 cohorts was indeed associated with shorter PFS and OS. Given that SCOTROC4 patients only received first line platinum monotherapy, this would suggest that higher RAD51 expression predicts for platinum-resistance, thus resulting in a shorter PFS and OS. In recurrent EOC, the presence of mutations in HRD related genes (40,41) and a PFI of more than 6 months better predicts responses to platinum-based chemotherapy, but currently no biomarkers exist to predict platinum resistance. Our results suggest that in the context of HRD negativity in EOC, high basal RAD51 expression may be such a resistance marker and will need further validation in previously concluded first-line trials of platinum therapy.

A crucial corollary of these observations will be whether high RAD51 protein levels also define a subset of patients with HR proficient EOC who will not benefit from PARPi maintenance therapy. Understanding the mechanisms and biomarkers of resistance to HR directed therapies are crucial to identify patients for whom alternative therapeutic options need to be considered or developed. Three PARPi (niraparib, rucaparib and olaparib) have recently been approved for the treatment of ovarian cancer patients. Current predictive markers of improved outcome following PARPi therapy (42) include HRD (43), prior platinum sensitivity(33) and genomic scar assays such as the Myriad myChoice® or Foundation Medicine LOH assays (33,44-47). Data from first line PARPi maintenance studies like PAOLA-1 (olaparib+bevacizumab vs. placebo+bevacizumab) and PRIMA (niraparib vs. placebo) showed that *BRCA* mutant and HRD positive tumours appear to benefit from PARPi maintenance therapy following first line platinum-based chemotherapy in advanced ovarian cancer. However, in the earlier NOVA (33) and ARIEL3 (48) trials, PARPi treatment conferred a survival benefit even in HRD negative but platinum sensitive EOC tumours. These clinical studies have highlighted two key points: firstly, while benefit from PARPi is greatest in tumoirs with BRCA1/2 mutations, followed by HRD positive tumours, the absence of either (based on currently validated genomic scar assays) does not exclude the possibility of benefit from PARPi maintenance; and secondly, platinum-response (i.e. RECIST partial or complete response) is currently a more robust surrogate marker of PARPi benefit than available scar assays (42). There remains a pressing need to improve patient selection strategies, particularly to identify those patients in whom no benefit is expected from PARPi therapy.

Given the aforementioned role of platinum-sensitivity as a key surrogate marker of PARPi benefit, the SCOTROC4 study represents a unique dataset in that all patients only received carboplatin monotherapy in the first line setting, and thus represents an optimal cohort of EOCs to interrogate predictive biomarkers of HR-directed therapy. Notably, in a subgroup analysis of SCOTROC4, we observed that the predictive value of RAD51_Vs_ for platinum resistance was restricted to the HRD negative (i.e. HR proficient) tumours. Why RAD51 expression does not appear to be predictive in HRD positive tumours is unclear. Possible explanations include that upregulation of RAD51 *per se* does not overcome a marked HR deficiency *in vivo*, or that the numbers of HRD positive tumours in the study are too small to discern any difference in outcome in this subgroup. Putting this observation together with analysis of genomic scars from the abovementioned PARPi trials suggests that a subset of HRD negative cases with high RAD51 expression are likely to be those that do not benefit from any HR-directed therapy (platinum or PARPi). This will warrant further exploration in tumour samples from the already completed phase III studies of PARP inhibitors in EOC (48,49).

To explore the functional impact of RAD51 overexpression in ovarian cancer, we developed a RAD51 overexpression model in HGSOC cell lines but did not observe resistance to platinum or olaparib in-vitro. It is well known however, that a significant determinant of cytotoxic efficacy in vivo is the immune and stromal microenvironment of tumours. Kroeger *et al.(50)* assessed co-localization patterns of immune cells in the high grade serous ovarian cancers and found that tumours containing CD8+, CD4+, and CD20+ TIL, together with APCs, were associated with markedly increased survival. When we applied spectral imaging to evaluate immune-infiltration, we found a statistically significant reduction of infiltrating CD3+/CD8+ cytotoxic T-cells in RAD51-High tumours (Fig.5) with no marked change in CD3+/FOXP3+ regulatory T-cell fraction (SuppFig.8). Notably, the immune exclusion phenotype we observed was primarily noted in tumours that are RAD51-High and *BRCA* wild-type and could account for the poorer outcomes observed in HR-proficient tumours. However, analysis of immune gene expression from multiple independent datasets noted that RAD51-High tumours show enrichment of cytokines of the CXCL9/10/11 family, which typically function to recruit T-cells into tumours. Reconciling these findings with the exclusion of cytotoxic T-cells in RAD51-High tumours suggests the existence of other “immune checkpoints” or hitherto undescribed negative regulatory features within these cases that counteract the CXCL family of cytokines. One attractive candidate is CD47, a “don’t-eat-me” signal for macrophages, which is consistently high in RAD51-High tumours across all four cohorts tested. An anti-CD47 antibody has shown promise in lymphomas (51,52), and could be evaluated in the context of these ovarian cancers as well. Future work in our laboratory will focus on the exact relationship between high RAD51 and possible negative immune regulators in cancer.

Taken together our findings suggest a possibility that high RAD51 expression constitutes a novel cell-intrinsic feature associated with evasion of immune surveillance in EOC (Figure 6). Aberrant DNA repair is known to generate immunogenic intermediates that signal to the immune system through cytosolic pathways for DNA sensing (53,54). To our knowledge, this is the first study to demonstrate the association between overexpression of a DNA repair protein and immune exclusion. This study is mainly hypothesis generating, being limited by the fact that it is a retrospective analysis of a subset of patients and should be further validated in prospective cohorts of clinical trials of PARPi or platinum-based chemotherapy. Nonetheless, given that high RAD51 expression appears to define a distinct subtype of HR proficient EOC for which platinum-based treatment may be less effective, the stage is set to develop better therapeutic approaches for cancers with this phenotype. Such approaches may include the addition of anti-angiogenic agents, e.g. bevacizumab, that have been shown to remodel tumour vasculature and reprogram the tumour microenvironment to facilitate anti-tumour immunity and enhance chemosensitivity (55,56). Comparing RAD51 protein with mRNA expression in large cohorts will also help identify which of these could serve as the most clinically applicable predictive biomarker or stratification factor in future clinical trials.

**Figure 6.**
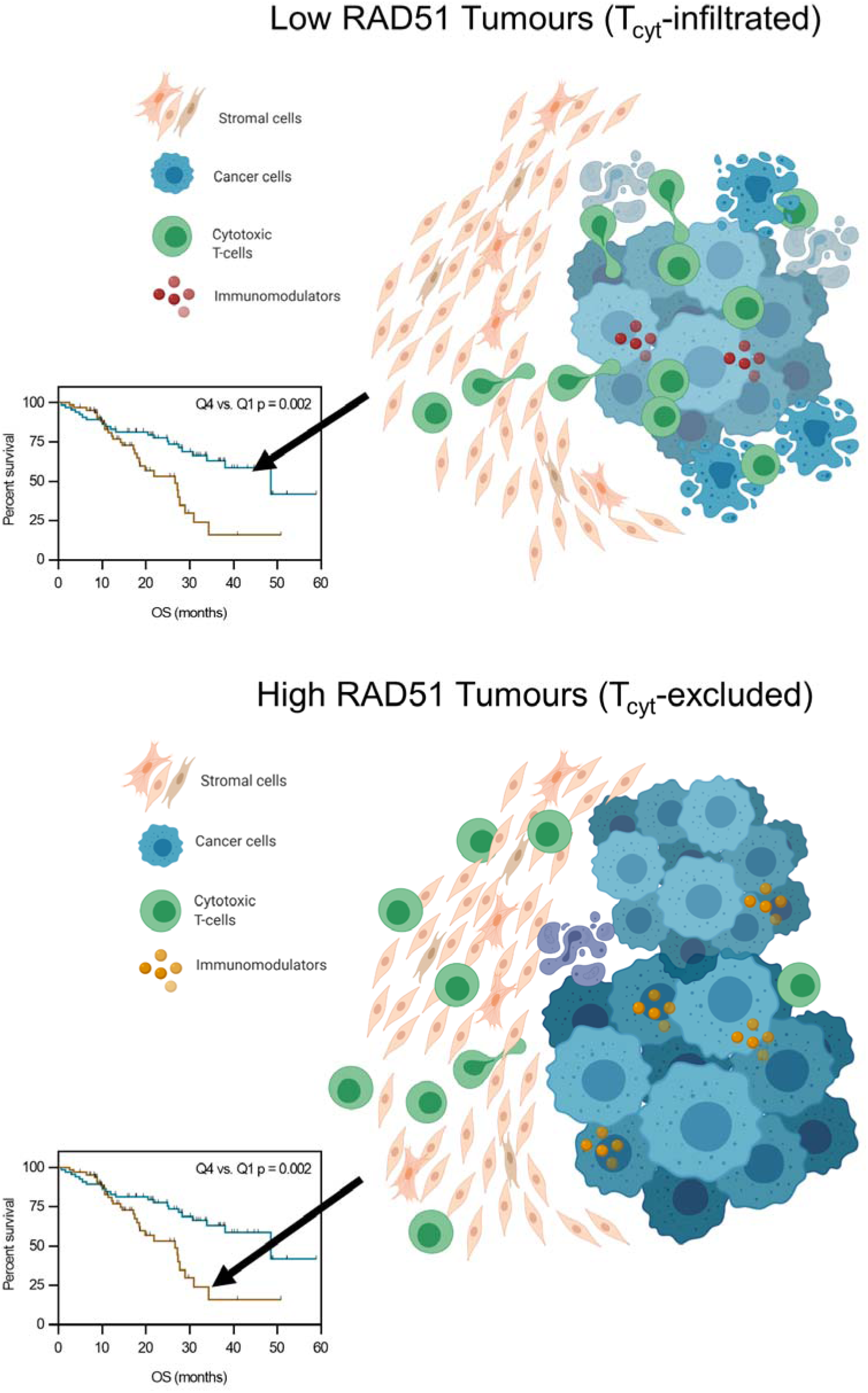
Schematic representation of the immune microenvironment in RAD51-High and -Low tumours.

## Acknowledgements

We thank the patients from the cohorts described in this paper, who generously consent to their tissues being analyzed for research.

## Author contributions

Experimental design: MMH, DSPT and ADJ; Multiplex IHC experiments: MMH; Pathological review: DL and BP; Bioinformatic analysis: JJP, TTZ, SL, ASS, KS, EC; Biostatistics: MMH, MH, HC, DSL, SL; Coordination of sample provision/ ethics approvals: RB, NP, DL, AK, DGH, DSPT, ADJ; Writing of the manuscript: MMH, JDW, AK, DGH, SK, RB, JJP, DSPT, ADJ.

## Data and materials availability

All data associated with this study are available in the main text or the supplementary materials. [3986/5000]

