## Supplementary methods and figures for "Primary platinum resistance and immune exclusion in ovarian carcinomas with high expression of the homologous recombination mediator RAD51"

Hoppe *et al.*

### Supplementary Methods

#### Fluorescent immunohistochemistry (fIHC) analysis

RAD51 is a key regulator of HR. Immunofluorescence of RAD51 shows clearly discernible microscopic foci, which are widely used as an *in vitro* surrogate of HR proficiency, and have also been tested in clinical material (1-3). RAD51 foci estimation as a quantitative readout is however hampered by wide variation in what is considered a “focus”, which is largely dependent on imaging parameters. In this study, we use the OPAL-TSA staining approach, which is not suited for subcellular localization, but is optimal for analysis of the per-cell expression of RAD51. Previous studies of RAD51 protein expression in cancer have also been limited by technical challenges in quantitation of data. Here we optimized a digital pathology platform – the Perkin Elmer Vectra, to quantitate RAD51 expression in clinical samples of EOC. We take into account the continuum of RAD51 staining, and use quartiles to study the correlation of RAD51 expression with biological markers and clinical phenotypes of interest.

Multiplexed fIHC was performed on formalin fixed paraffin-embedded (FFPE) whole-tissue sections or tissue microarrays (TMA) to assess protein expression levels of markers of interest. Opal 7-Color Manual IHC Kit (PerkinElmer Inc., Waltham, MA, USA) was used to visualize targets following the manufacturer’s protocol. Briefly, 3μm thick tissue sections were deparaffinized in organic solvents and rehydrated using a gradient of ethanol solutions. Heat-mediated antigen retrieval was performed on rehydrated slides using Target Retrieval solutions (Dako, Denmark), followed by primary antibody incubation. Primary antibodies used in this study, dilution and incubation time are listed in Supplementary table 3. Slides were then washed in 0.1% Tween20 (Sigma-Aldrich) solution in water 3 times, 2 minutes each wash with agitation. Secondary antibody incubation was 10 minutes at room temperature using anti-mouse or anti-rabbit IgG HRP-labelled secondary antibodies (1:1000, PerkinElmer Inc., Waltham, MA, USA) followed by a washing cycle. Finally, slides were incubated with choice of Opal fluorophore (1:100) for 5 minutes at room temperature, followed by a wash cycle. This constitutes a full cycle of antibody staining and can be multiplexed by repeating this sequence starting from heat-mediated antigen retrieval using a microwave to strip the specimen of antibodies present from the previous round. DAPI was added to the secondary antibody mixture to serve as a counterstain. Slides were mounted using Mowiol 4-88-based mounting medium (Sigma-Aldrich).

Slides were imaged using the Multispectral Vectra 2 Imaging System (PerkinElmer Inc., Waltham, MA, USA). Multispectral images were analyzed using the inForm 2.2 software (PerkinElmer Inc., Waltham, MA, USA). To obtain monochrome images of all components of a multiplexed fIHC slide, images were first unmixed using a prepared spectral library of fluorophore-specific pure spectra measured from single stained slides of each fluorophore. EOC-specific autofluorescence spectra was also obtained from an unstained, but processed slide. Next, a trainable tissue segmentation algorithm was used to identify regions of interest in images i.e. epithelial tumour cells, but not stromal cell. With each multiplexed fIHC, we included a staining against EpCAM, an epithelial marker. Correct tissue segmentation was reviewed to ensure reliability of the segmentation protocol. All cells included in regions positively stained with the anti-EpCAM antibody were included in the tumour-specific cell-based analysis. Cells were segmented using the cell segmentation algorithm, creating a nuclear mask for each cell within the tumour region (see Fig.1B). The nuclear mask was based on the counterstain and correct segmentation was aided by the presence of the EpCAM membrane marker to help define cellular boundaries. Mean nuclear fluorescent intensity of a marker of interest was expressed as a Vectra score and used as a read-out of protein expression. The Vectra score is expressed in Normalized Counts as follows:

$$NormalizedCounts=\frac{fluorescentcounts}{2^{bitdepth}\times exposuretime\times gain\times binningarea}$$

Exposure time is expressed in seconds and binning area is 1 for all images. TMA cores with less than 100 tumour cells, damaged tissue or unwarranted staining patters were not included in further analysis and were considered as failed quality control (QC). In the SCOTROC4 cohort, median number of cells analysed per TMA core was 1596 (range 124-4590) and at least 2 TMA cores were analysed per patient (range 2-12).

Routine biomarker evaluation centres on group stratification across the median value. However, as RAD51 Vectra score follows a normal distribution within the cohort (Fig.1A and Fig.3A), stratification across the median is non-optimal. Division across median dichotomizes the peak cases into -High and -Low groups, saturating them with cases of quantitatively similar results. To preserve the biological distinction relating to RAD51 expression in a normally distributed cohort, we reasoned that stratifying the cohort into three biologically distinct groups will reveal truer associations between RAD51 and survival. Thus, we divide cohorts into RAD51-Low cases within the first quartile (Q1), RAD51-High cases within the fourth quartile (Q4), and RAD51-IQR cases within the interquartile range (IQR, quartiles 2+3).

#### Creation and validation of RAD51 overexpressing cell lines

Four HGSOC cell lines were chosen to perform in-vitro experiments based on previous assessment (4): TYK-nu (*TP53*_mut._, *BRCA1/2*_wild-type_), Caov-3 (*TP53*_mut._, *BRCA1/2*_wild-type_), COV362 (*TP53*_mut_, *BRCA1*_mut._*, BRCA2*_wild-type_), FUOV-1 (*TP53*_mut._, *BRCA1/2*_wild-type_). The coding sequence (CDS) of the canonical RAD51 transcript (NM_002875.4) was amplified from normal fallopian tube cells (FT33) using AAAAGGATCCGCAATGCAGATGCAGCTTGA (forward) and AAAAGCGGCCGCTCAGTCTTTGGCATCTCCCA (reverse) primers and cloned into the BamHI and NotI cloning sites of the pMSCV-puro-Flag retroviral vector. HGSOC cell lines were transduced with virus containing the exogenous Flag-RAD51 CDS or empty vector and cultured in DMEM medium supplemented with 25mM HEPES, 10% FBS and 1μg/ml puromycin; expression of exogenous RAD51 was confirmed by western blotting (SuppFig.3A).

To confirm functionality of the Flag-tagged exogenous RAD51 construct, cell survival assays were performed. HGSOC cells lines were seeded at low confluency on a 96-well dish in three technical triplicates, 12h later siRNA transfection was performed. Lipofectamine RNAiMAX (ThermoScientific) transfection mixture was prepared according to manufactures protocol and RNA was added at 20nM; following target sequences were used: siControl - ON-TARGETplus Non-targeting Pool (Dharmacon), siRAD51-CDS – CAGAUUGUAUCUGAGGAAA, siRAD51-3'UTR - TCTTCCTGTTGTGACTGCCAGGATA. Twenty-four hours later cell culture media was changed and carboplatin (Sigma) was added at 1mM concentration and serially diluted. Ninety-six hours after drug addition, MTT solution was added and cells were kept for an extra 2h at 37°C to allow crystals for form. Next, Stop solution (37% (v/v) N,N-dimethylformamide, 14.2% (m/v) SDS and 2% acetic acid) was added and the plate was incubated on a shaker to allow complete dissolution of violet crystals. Absorbance at 564nm was read for each well by a Tecan Infinite 200 PRO plate reader. The extend of light absorbance was considered to be proportional to cell density.

For cell-survival assays shown in Figure 5 and Supplementary figure 4, untransfected cells were treated as described above with carboplatin or olaparib (AZD2281, Selleck).

#### GSEA and differential gene-expression analysis

Total RNA was isolated form RAD51-overexpression and control HGSOC cell lines using RNeasy Mini Kit (Qiagen) and standard polyA sequencing was done by NovogeneAIT. Identification of differentially expressed genes and Gene Set Enrichment Analysis (using GO geneset library) comparing RAD51-overexpression vs. control cell lines was performed using standard pipelines available publicly on the CSI NGS Portal (5).

To identify differentially expressed immune genes between RAD51-High (fourth quartile, Q4) and -Low (first quartile, Q1) cases, the processed high-grade endometrioid, high-grade serous ovarian cancer gene expression data of TCGA (*n=*566) (6), Australian Ovarian Cancer Study (AOCS, GSE9891; *n*=267)(7), Massachusetts General Hospital (MGH, GSE26712; *n*=185) (8) and Duke University Hospital (Duke, GSE3149; *n*=146) (9) were extracted from CSIOVDB (10). Quartiles were calculated independently for each dataset. Statistical analyses were conducted using Matlab® R2016b version 9.1.0.960167, statistics and machine learning toolbox version 11.0 (MathWorks; Natick, MA, USA); t-test was applied for identifying differentially expressed genes.

### Supplementary Figures

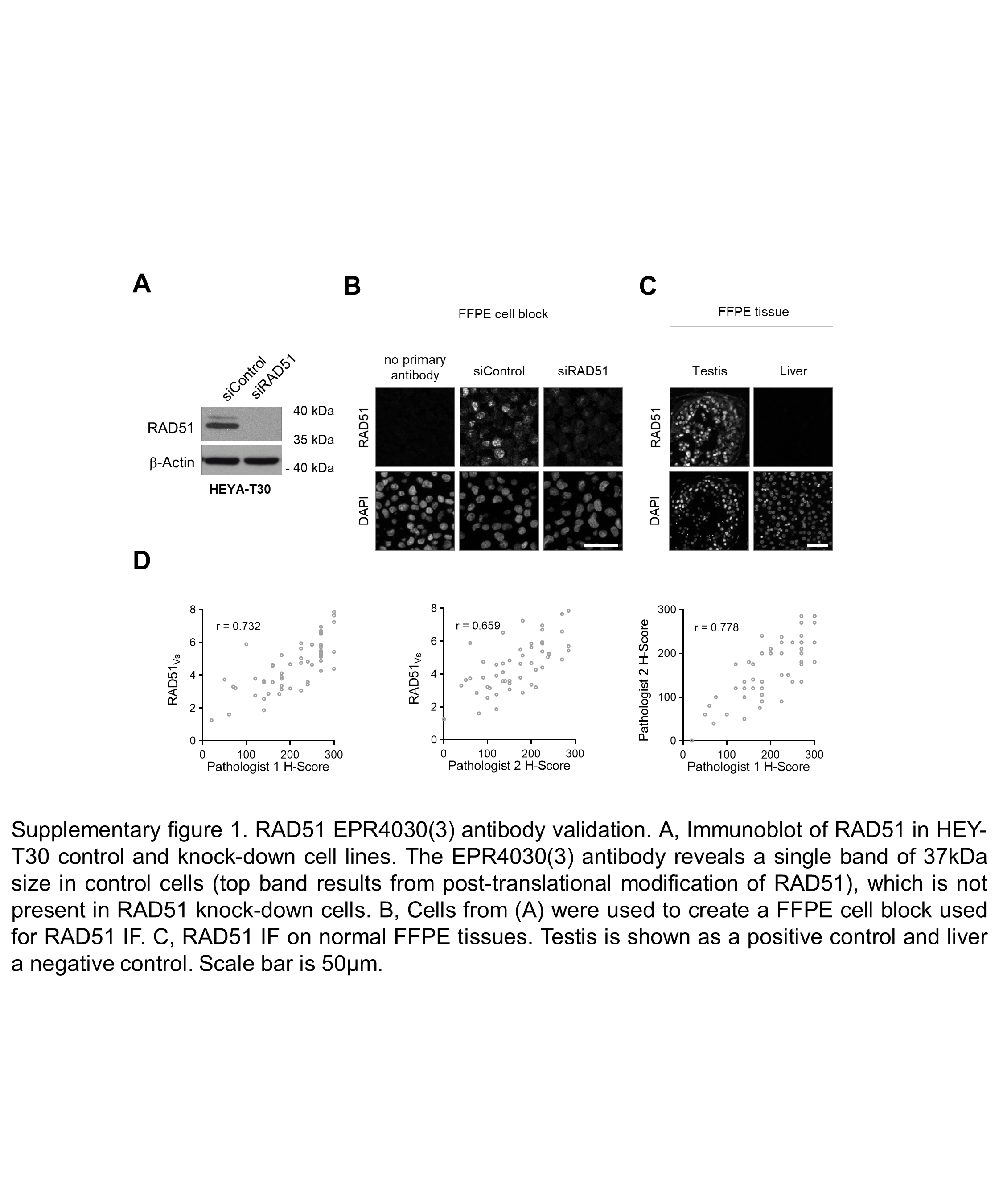

**Supplementary figure 1**. RAD51 EPR4030(3) antibody validation. **A**, Immunoblot of RAD51 in HEY-T30 control and knock-down cell lines. The EPR4030(3) antibody reveals a single band of 37kDa size in control cells (top band results from post-translational modification of RAD51), which is not present in RAD51 knock-down cells. **B**, Cells from (A) were used to create a FFPE cell block used for RAD51 IF. **C**, RAD51 IF on normal FFPE tissues. Testis is shown as a positive control and liver a negative control. Scale bar is 50μm. **D**, Correlation in a cohort of EOC cases of RAD51_vs_ and two independent pathologist H scores (Left and middle panels) and correlation of two pathologist H scores (right panel).

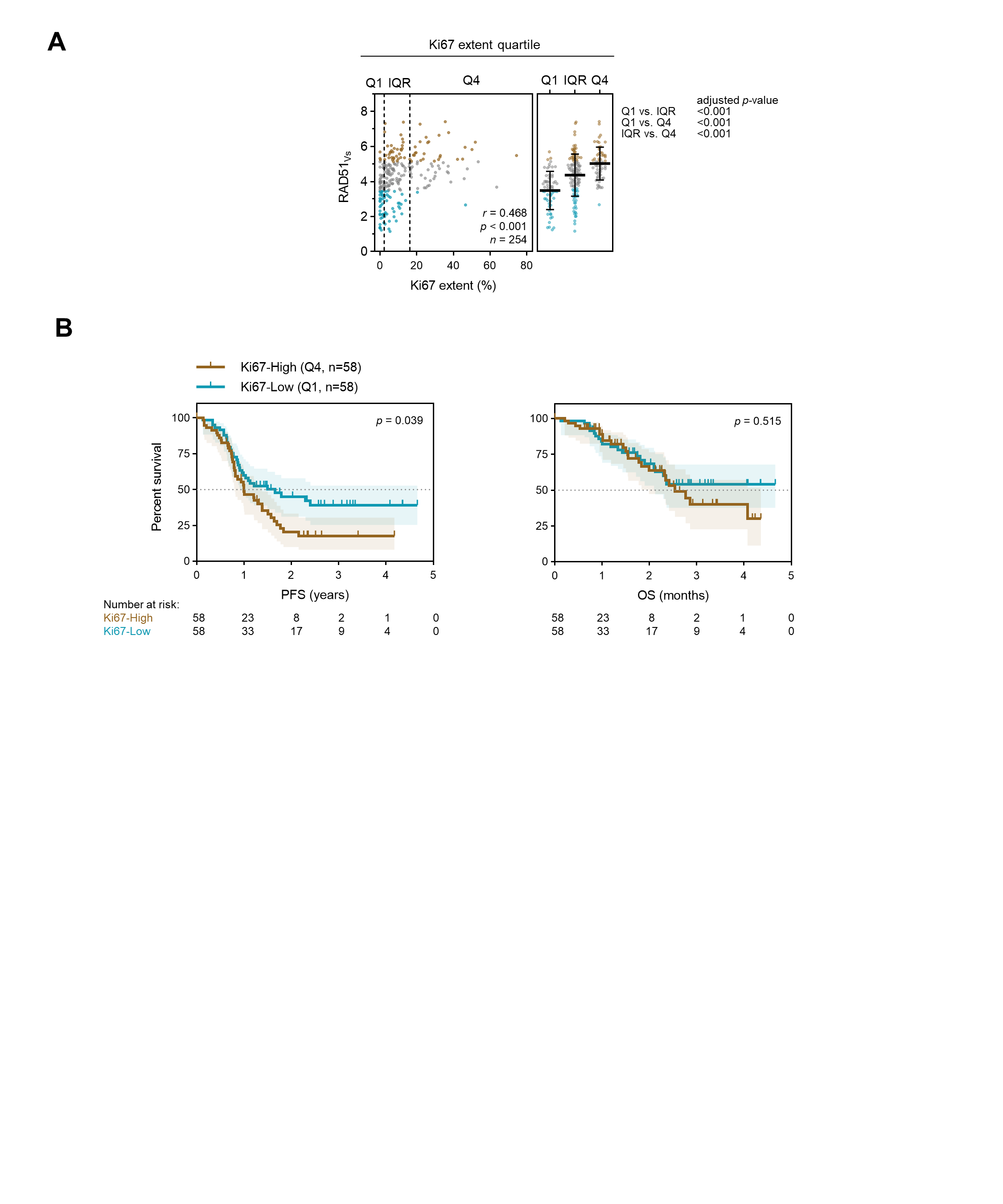

**Supplementary figure 2.** Proliferation analysis of the SCOTROC 4 cohort. **A**, Correlation of Ki67 extent and RAD51_Vs_. Spearman correlation (*left*) and one-way ANOVA with Bonferroni correction (*right*). Mean with standard deviation. **C**, Survival analysis of the SCOTROC4 cohort. Kaplan-Meier plots for PFS (*left*) and OS (*right*) stratified according to quartiles of Ki67 extent. Log-rank test.

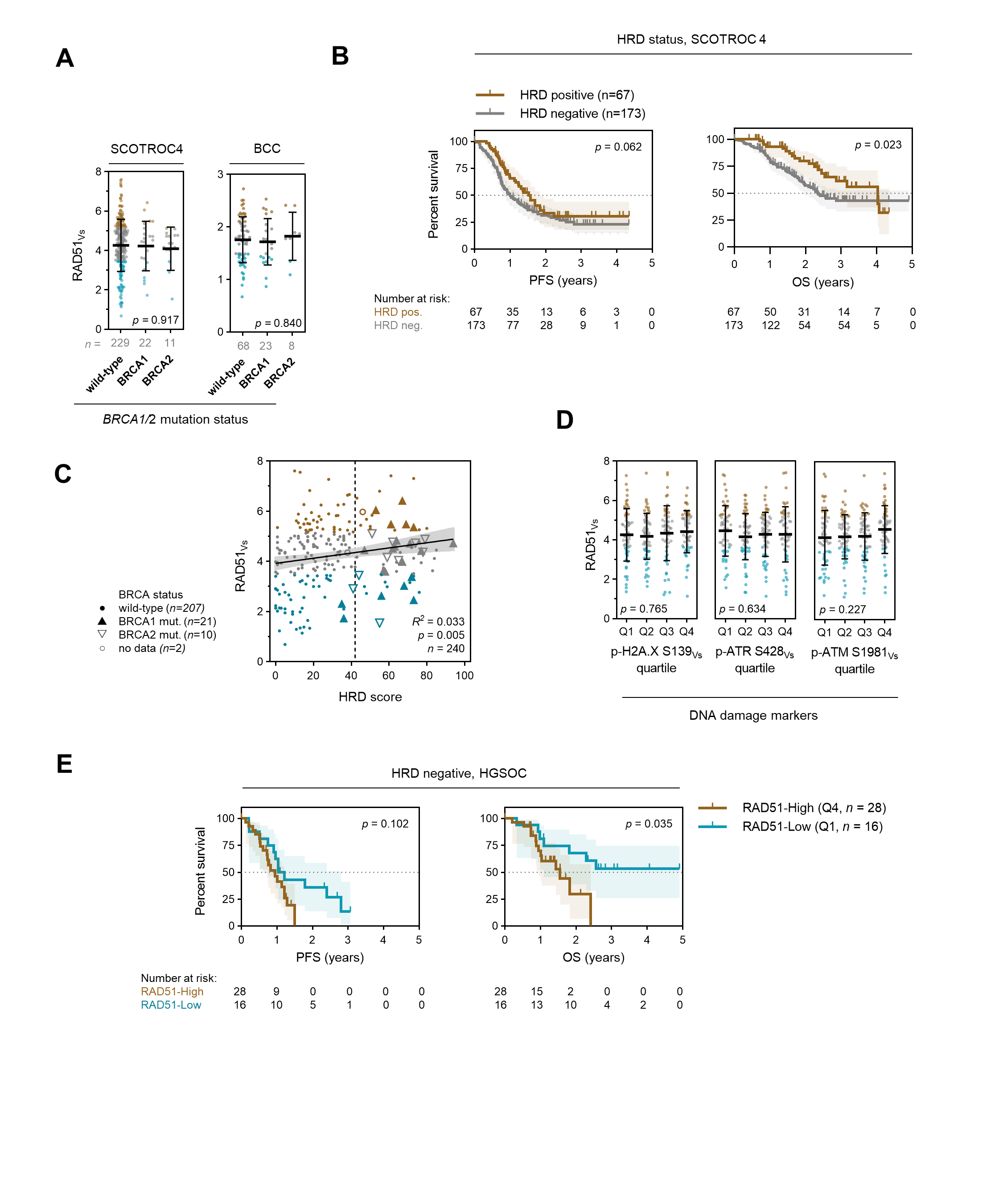

**Supplementary figure 3.** No correlation of RAD51_Vs_ and genomic instability burden **A**, Correlation of RAD51_Vs_ with *BRCA* mutation status in EOC. One-way ANOVA. **B**, Kaplan-Meier plots for progression-free survival (PFS) (*left*) and overall survival (OS) (*right*) stratified according to HRD status. **C**, Linear regression of 'genomic scar’ HRD score assay and RAD51_Vs_. Vertical dashed line denotes HRD positivity score of 42. **D**, Correlation of RAD51_Vs_ with Vectra scores of DNA damage markers. Q – quartile. One-way ANOVA. Mean with standard deviation. **E**, HRD negative HGSOC cases stratified according to quartile of RAD51_Vs_. Log-rank test.

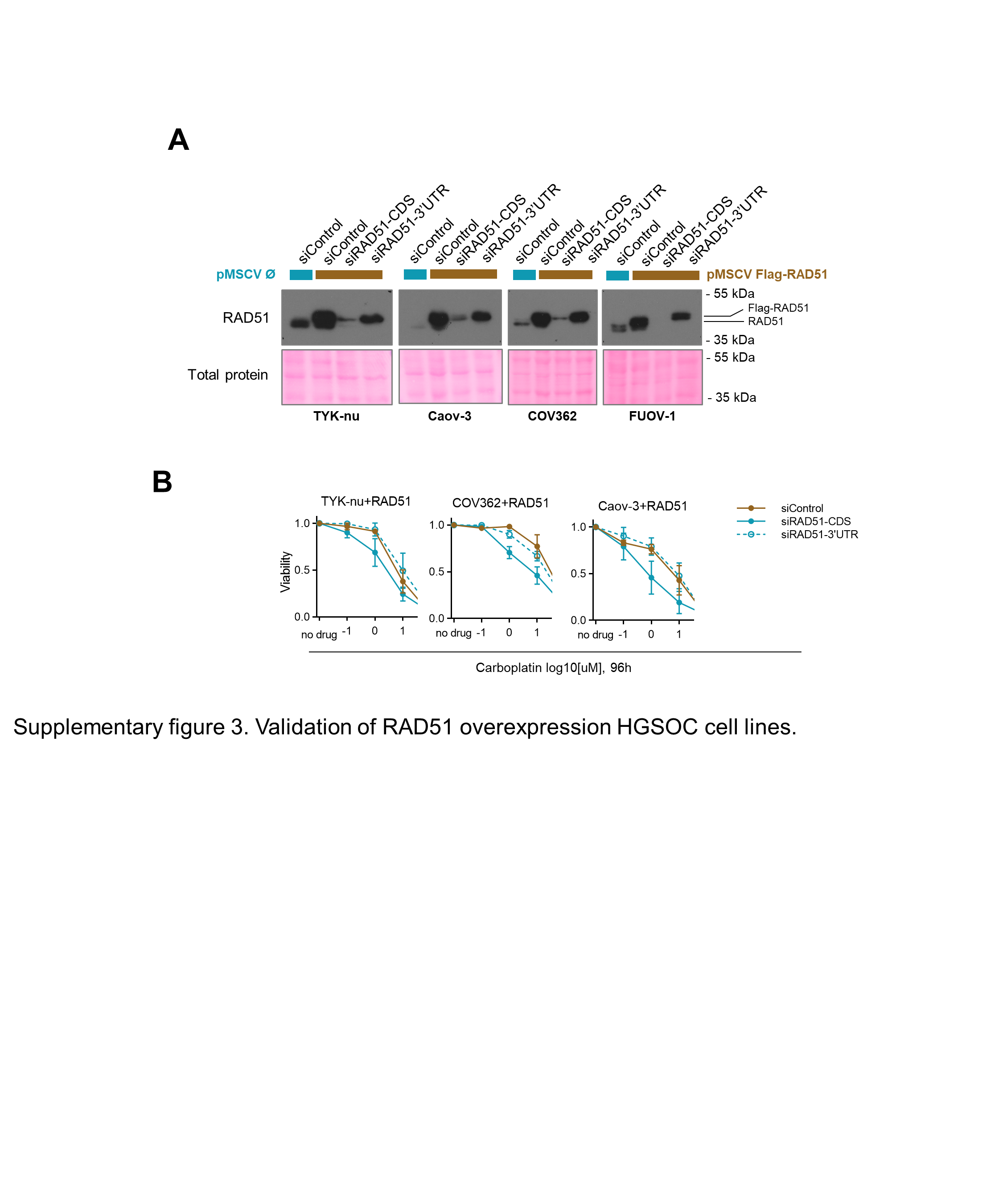

**Supplementary figure 4.** Validation of RAD51 overexpression in HGSOC cell lines. **A**, Immunoblot of RAD51 upon overexpression and subsequent RNAi-mediated depletion of total or endogenous RAD51 mRNA. **B**, Functional validation of exogenous Flag-RAD51. Flag‑RAD51 rescues carboplatin sensitivity upon depletion of endogenous RAD51 protein.

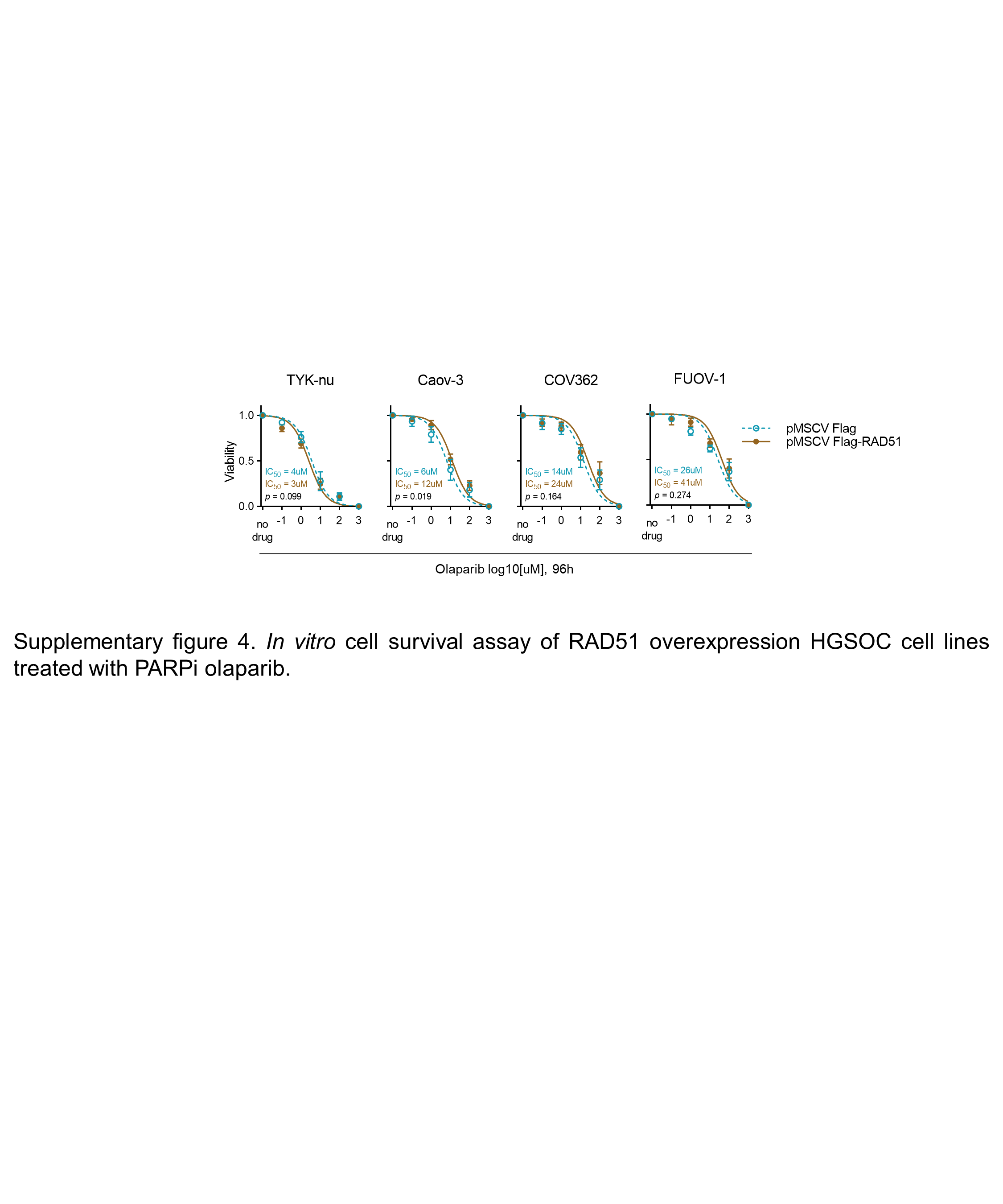

**Supplementary figure 5.** *In vitro* cell survival assay of RAD51 overexpression HGSOC cell lines treated with PARPi olaparib.

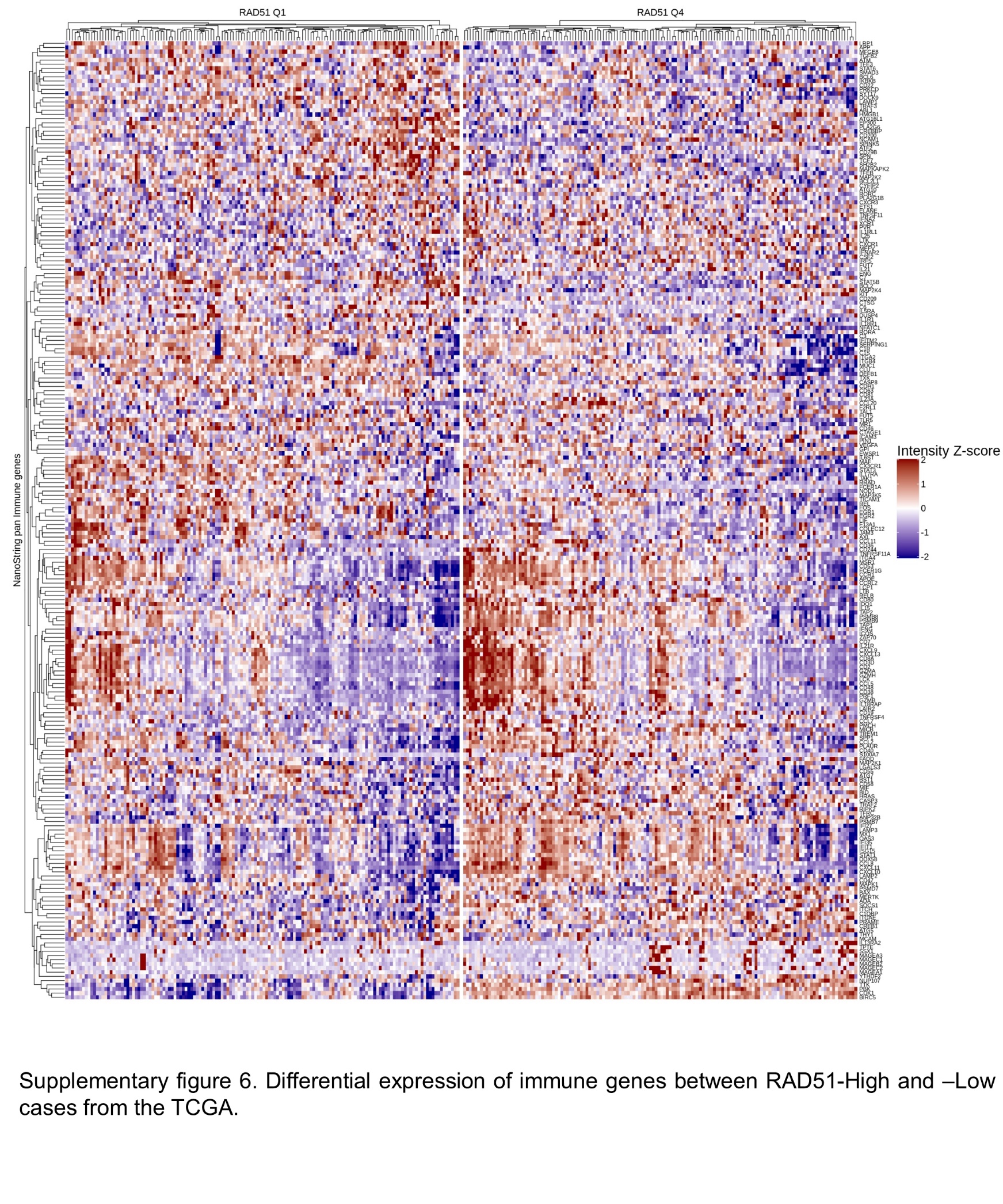

**Supplementary figure 6.** Differential expression of immune genes between RAD51-High and –Low cases from the TCGA.

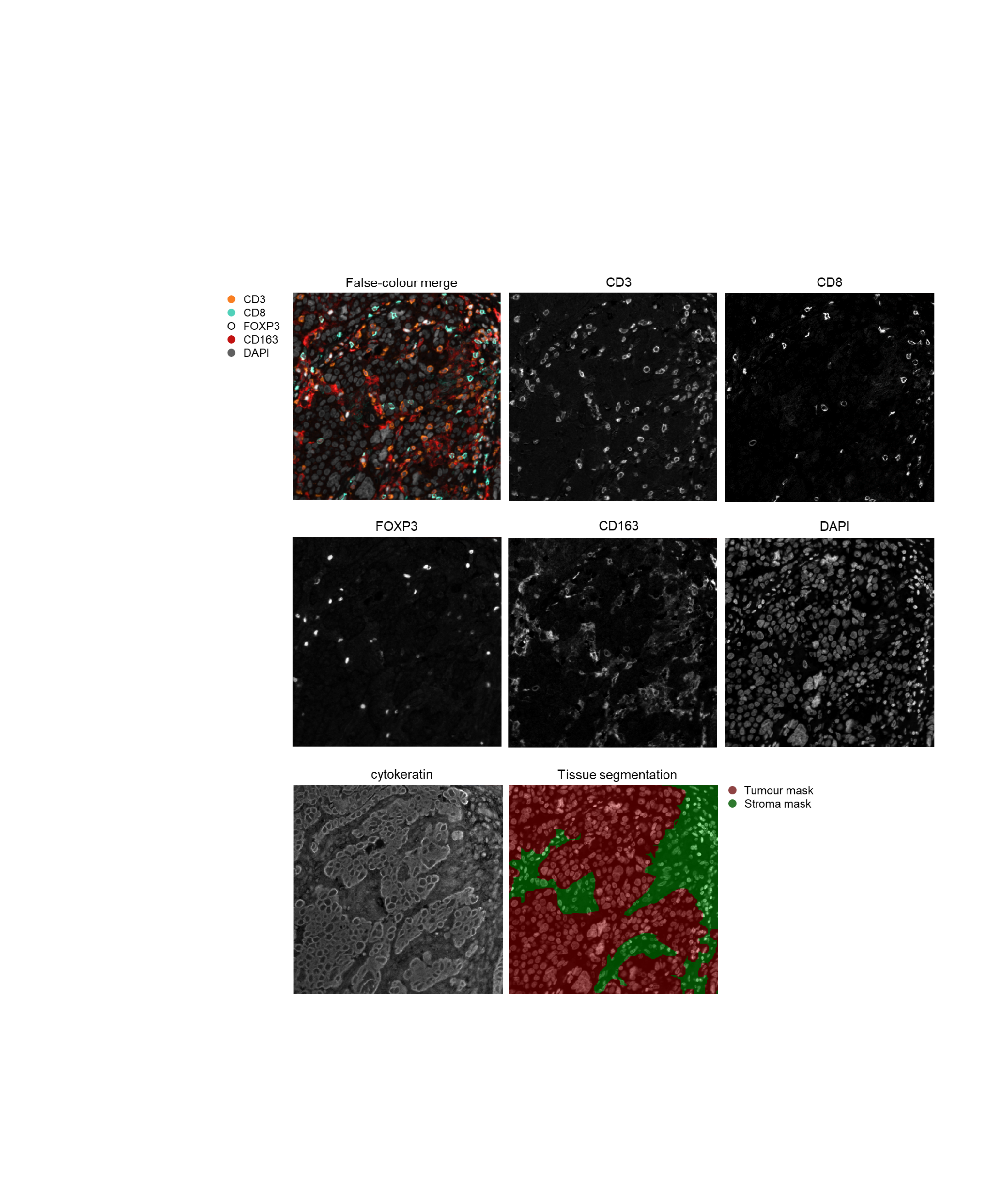

**Supplementary figure 7.** Immune infiltration analysis by multiplexed fIHC. Unmixed monochrome components are shown along with a false-coloured merge image. Cytokeratin staining was used to differentiate between the tumour and stromal compartments of the sample.

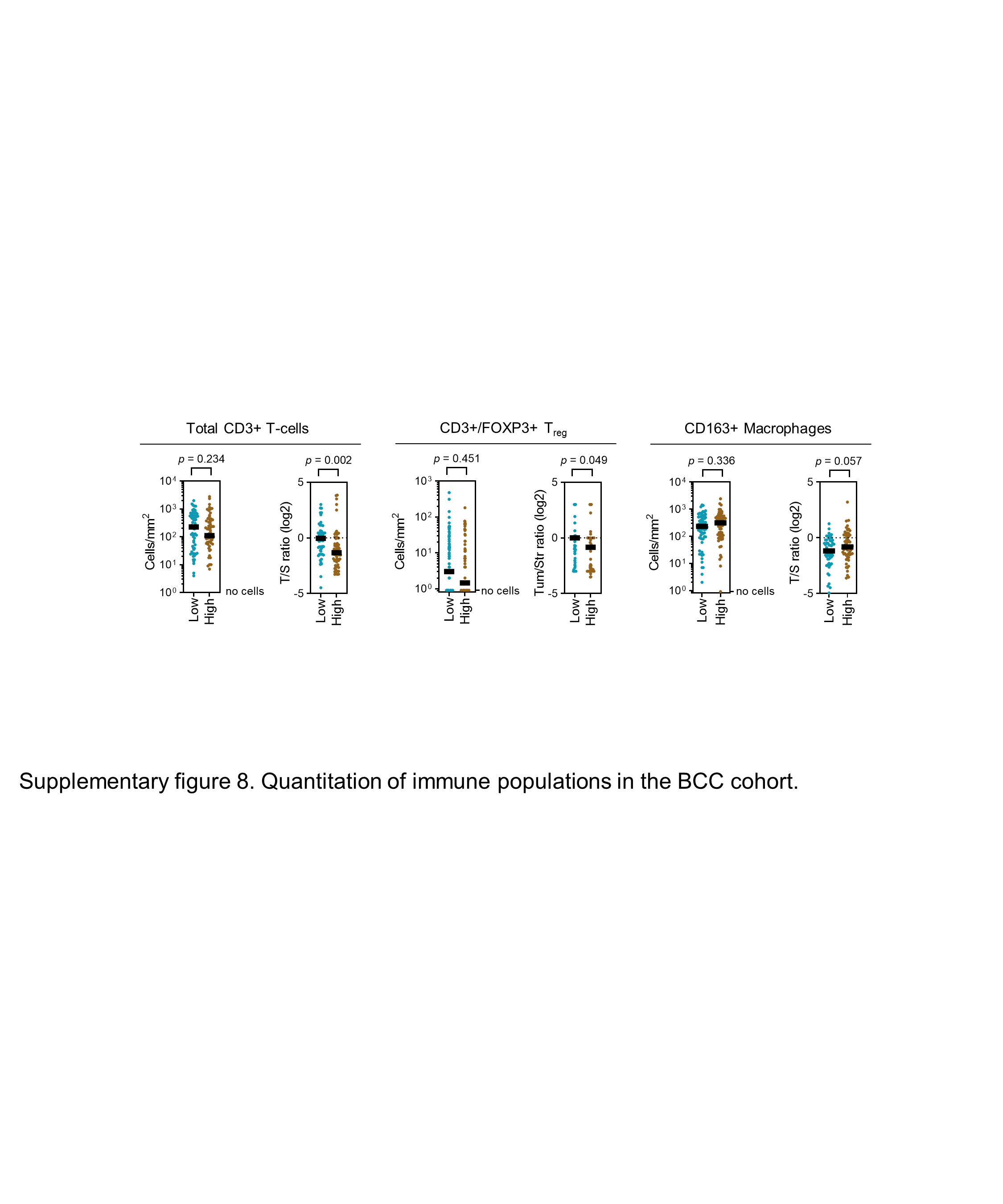

**Supplementary figure 8**. Quantitation of immune populations in the BCC cohort. Results for RAD51-High and -Low tumours are shown. Mann Whitney test. T/S – tumour/stroma ratio.
